# The first proof of concept demonstration of nanowarming in coral tissue

**DOI:** 10.1101/2023.03.16.533048

**Authors:** Jonathan Daly, Jessica Bouwmeester, Riley Perry, Chris Page, Kanav Khosla, Joseph Kangas, Claire Lager, Katherine Hardy, John C. Bischof, Mary Hagedorn

## Abstract

Climate change is causing ocean acidification and warming, resulting in mass bleaching and death of corals globally. Cryopreservation and biobanking to secure the genetics of threatened populations is currently limited to coral sperm and larvae, which are only available during brief annual spawning events and are impacted by ocean warming, so there is an urgent need for methods to enable biobanking activities year-round. Here, we used vitrification and nanowarming to successfully recover adult coral tissues in a novel sample type, the single-polyp microfragment (SPMF). Fluorescence and confocal microscopy showed clearly defined tissues and green fluorescent protein fluorescence around the polyp mouth post-warming in 43.3% of SPMFs at 24 hours post-warming, and 30.0% at one month. These advances provide a basis for continued research and development of a field-ready protocol for cryopreservation of adult coral tissues, to permit biobanking of threatened coral species throughout the year and support reef restoration efforts.

## Introduction

Repeated ocean warming events over recent years have resulted in the stress and death of large numbers of coral colonies on reefs worldwide^1^, and there is the very real possibility of losing all but a few reefs globally by mid-century^2 3 4^. In addition to the loss of individual coral colonies and the structure that they provide on the reef, there is also the loss of potential future generations of coral that the affected colonies would have produced. The genetic information of these corals and their descendants is therefore lost from the population, diminishing the overall genetic diversity, and potentially impacting the ability of reefs to adapt to warming oceans. By using cryopreservation to secure the genetics of living coral, we can help to ensure that the genetic information and future offspring of those colonies are not lost completely^5^. Secure storage in biorepositories can ensure that these cryopreserved samples remain viable for decades and can be incorporated into coral aquaculture production systems to supply restoration, or used to support adaptation research^6^.

Cryopreservation of coral sperm began in 2006^7^ and there are currently thousands of cryopreserved samples in biorepositories worldwide. However, the use of cryopreserved coral sperm requires access to coral eggs, meaning that both sperm cryopreservation and its use are tied to coral reproduction. Coral reproduction is highly seasonal, with most species only spawning for a few nights per year following the full or new moons in Spring or Summer. Most corals are hermaphrodites that spawn by releasing gamete bundles into the water that break up to release the sperm and eggs^8^. Gametes typically remain viable for a short period (<5 hours) after bundle break up, meaning that the cryopreservation of sperm, and its use for reproduction, are limited to a window of only a few hours per year. Adding to these challenges, reproduction and in particular sperm motility are negatively impacted by ocean warming events^9 10^, resulting in diminished post-thaw outcomes. The recent development of larval cryopreservation^11^ expands the period during which coral samples are available for cryopreservation to a few days per year, but is still reliant on reproduction. An alternative approach to securing the biodiversity of corals would be to cryopreserve adult coral tissues in the form of microfragments, which are produced by cutting fragments of adult coral colonies into smaller pieces, typically >1cm^2^. Microfragmentation takes advantage of the natural ability of corals to reproduce asexually and is a common propagation tool for reef restoration^12^.

For cryopreservation and biobanking, adult coral tissues have the advantage of being available all year round; however, they have the disadvantage of being highly complex, containing algal symbionts and a calcium carbonate skeleton, and of being sensitive to chilling injury^13^. One of the most effective tools available for the cryopreservation and recovery of complex tissues is vitrification and laser warming with the aid of plasmonic gold nanorods, referred to as ‘nanowarming’. Laser nanowarming technologies^14^, previously used successfully with coral larvae^11^, permit ultra-rapid warming of vitrified microlitre volumes at rates up to 10^7^ C/min^15 16^, enabling the use of lower-concentration vitrification solutions that are less toxic to cells and tissues. For smaller sample volumes (e.g., mouse embryos (80 µm diameter) in 0.1µL droplets) the use of extracellular laser absorbers in the vitrification solution is sufficient to exceed the critical warming rate via the diffusive transfer of heat to the centre of the sample^17^. However, in the case of larger samples like zebrafish embryos (1 mm diameter) the application of external warming alone creates a large thermal gradient as heat attenuates across the tissues, resulting in insufficient warming at the centre of the sample and subsequent ice nucleation. To overcome this, Khosla et al. ^14 18^ microinjected plasmonic gold nanorods that absorb at 1064 nm along with cryoprotectant into zebrafish embryos while also deploying around the embryos to provide both internal and external warming, resulting in the successful recovery of living embryos, that could survive and develop to reproductive maturity. The maximum size of a particular tissue type that can be successfully recovered using vitrification and laser nanowarming is therefore determined by the physical limitations of heat transfer across the tissues and whether nanoparticles can be applied in both internal and external compartments. Additionally, the laser system typically used for nanowarming (the LaserStar iWeld used in the present study) has a maximum beam diameter of 2 mm, meaning that the biological sample and most of the surrounding droplet of vitrification solution should fit within this range to ensure adequate warming across the sample. Therefore, the physical limitations of the laser used for these studies sets strict boundaries for the tissue parameters in terms of their overall size and thickness.

The goal of the present study was to assess the utility of vitrification and laser nanowarming for cryopreservation and recovery of adult coral tissues. To achieve this, we first developed a method for producing coral microfragments that were small enough to fit within the physical parameters of the laser warming system. We then undertook a series of experiments to develop a vitrification solution along with equilibration and rehydration protocols for these single-polyp coral microfragments (SPMFs). Finally, we assessed the recovery of SPMFs following vitrification and laser nanowarming. The results of this study show for the first time that cryopreservation of adult coral tissues could be possible as a tool for securing coral reef biodiversity and provide a basis for future development of biobanking methods for coral fragments.

## Results and Discussion

In the present study, the Hawaiian coral *Porites compressa* was used to generate material for vitrification and laser warming experiments (Fig. 1). Most reef-building corals are colonies made up of thousands of coral polyps, each of which is an individual animal set inside a cup-like ‘corallite’ in the calcium carbonate skeleton that they secrete. The structure of the coral colony is predominantly composed of skeleton with the coral polyps and surrounding tissue forming a thin layer embedded in the surface of the skeleton. In *Porites compressa* the coral polyp and corallite together are approximately 1–1.5 mm in diameter; however, the conformation and mass of skeleton associated with the corallite mean that the coral in its natural form is not amenable to vitrification and laser warming, or to microinjection of cryoprotectants and nanoparticles. To generate material that would fit within the physical requirements of laser warming we used a technique called microfragmentation, which takes advantage of the natural ability of coral to reproduce asexually and to grow new tissue onto a suitable substrate. However, the highly specific physical parameters required for the laser nanowarming system necessitated a further refinement of this technique to produce a much smaller sample type with minimal skeleton, resulting in the production of SPMFs. The generation of this novel sample type also necessitated the development of an advanced coral husbandry approach to ensure a reliable supply of quality samples for experimentation.

**Figure 1.**
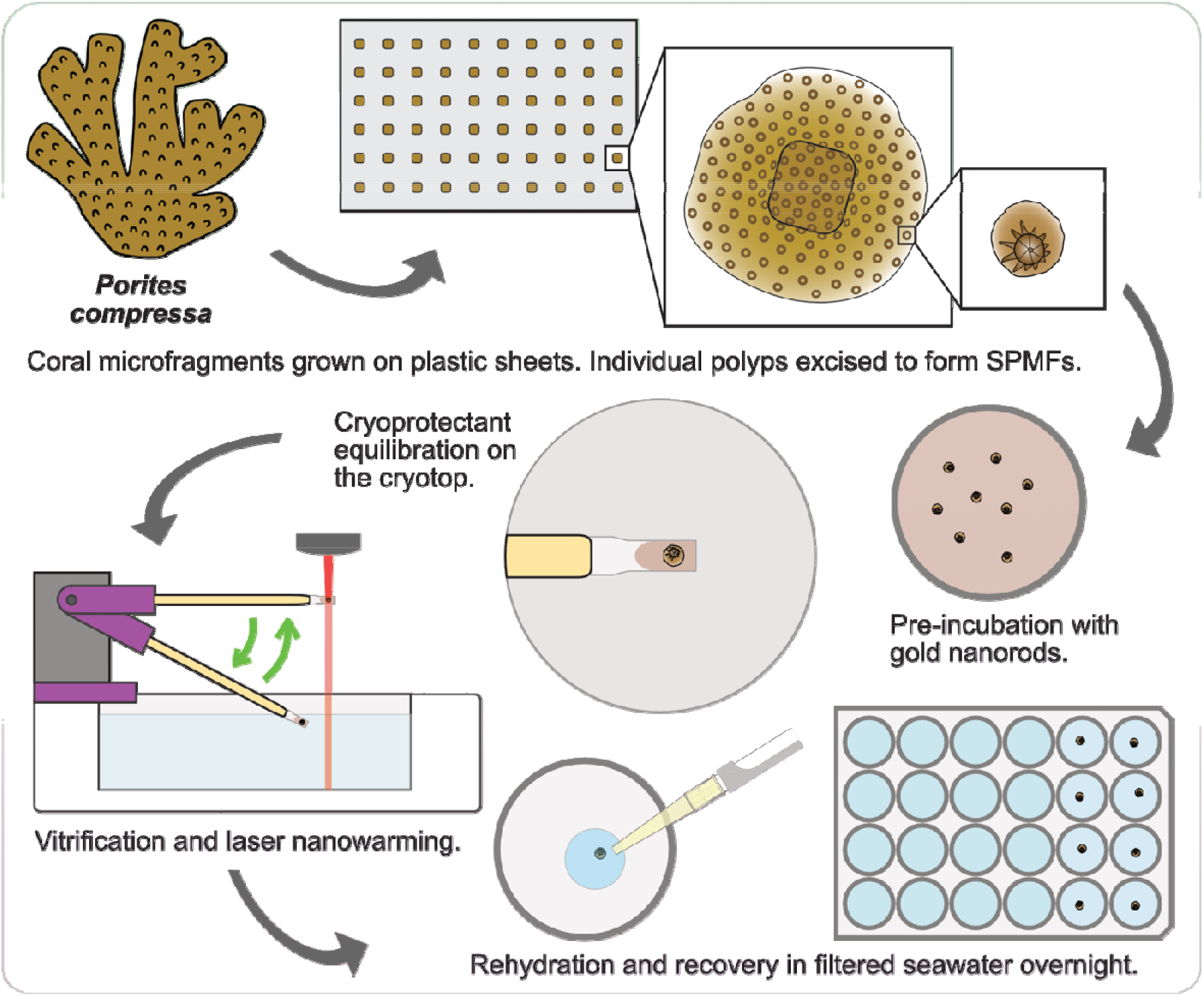
Production, cryopreservation, and recovery of single-polyp microfragments (SPMFs) from the coral *Porites compressa*. Corals were cut into microfragments and grown on plastic sheets, then individual polyps were excised from thin primary growth areas using a biopsy punch and allowed to heal in an incubator to form single-polyp microfragments (SPMFs). Single-polyp microfragments were pre-incubated in 0.07 mg/mL (4.5 × 10^11^ np/m^-3^) 1064 nm resonant PEG-coated gold nanorods in FSW for 1 hour, then equilibrated with cryoprotectant solution in three steps over 4 minutes on the cryotop. The cryotop with SPMF was transferred to the cryo device for vitrification and laser nanowarming using an iWeld 980 Series, 60-joule, Nd:YAG unpolarised infrared laser (240 V, 9.0 ms, ∼ 8.2 J). Single-polyp microfragments were then rehydrated in five steps over 4 minutes, followed by recovery in a 24-well plate overnight in an incubator without light.

### Generation of single-polyp microfragments

Optimised coral growth for SPMF production was achieved through strict maintenance of abiotic and biotic conditions, and several qualitative metrics were devised to avoid suboptimal grow-out conditions ^19^. Coral health metrics such as polyp extension, colour, and new tissue condition were regularly monitored, along with metrics gauging the health and progression of desirable fouling organisms (e.g., crustose coralline algae, CCA) to confirm stable tank conditions. A decline in value of these metrics often corresponded with suboptimal coral health and skeletal thickening, which rendered microfragments unusable for cryopreservation. Assuming optimal grow-out conditions, newly cut microfragments could produce thinly-sheeted new growth with rows of polyps suitable for creation of SPMFs (Fig. 2A) within four to six weeks.

**Figure 2:**
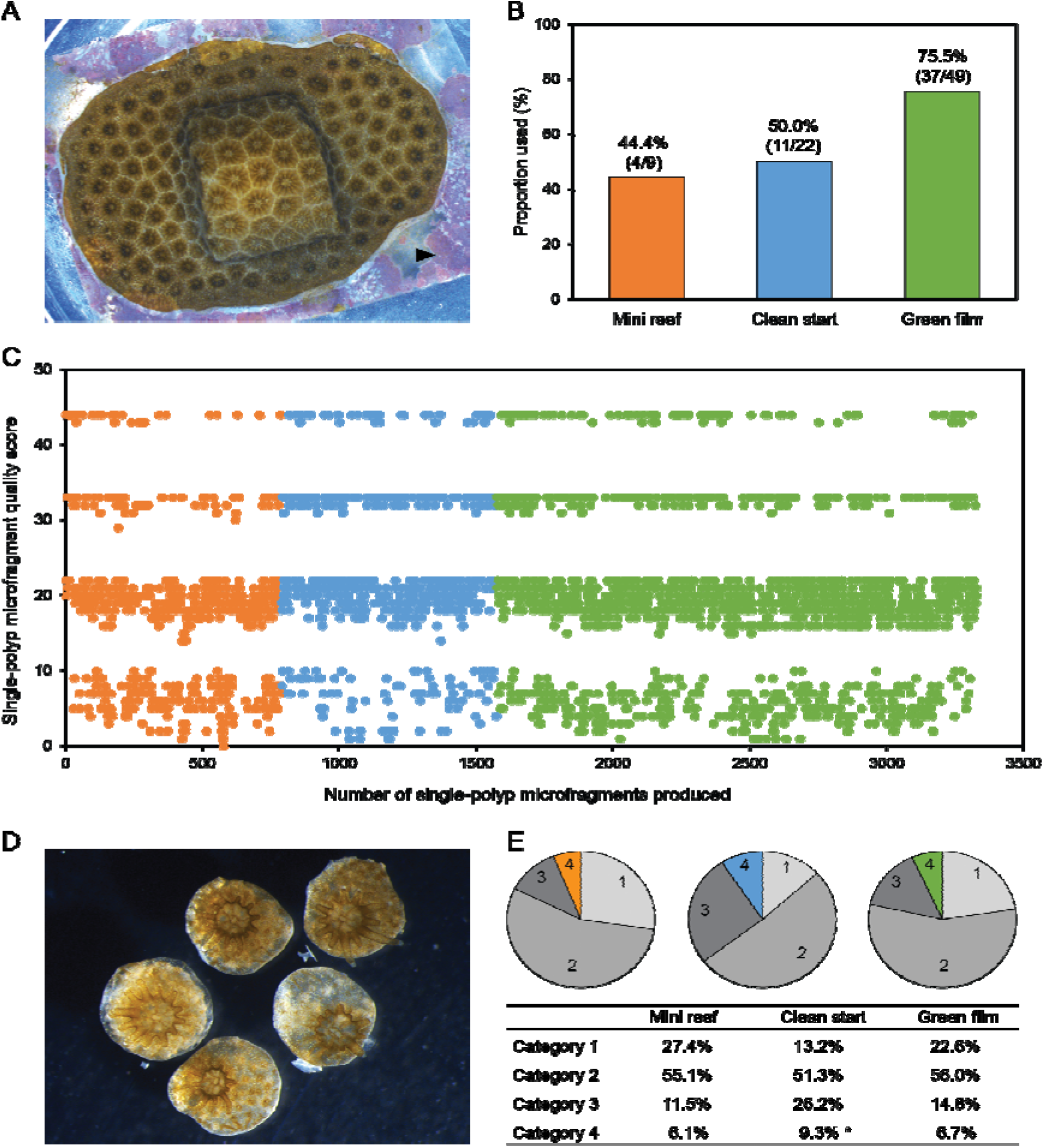
Production of single-polyp microfragments from the coral *Porites compressa* using three tank management methods: mini reef (MR; orange), clean start (CS; blue), and green film (GF; green). **(A)** New coral growth with rows of coral polyps surrounding the original microfragment and crustose coralline algae visible on the plastic sheet (arrowhead). **(B)** The proportion of *Porites compressa* colonies in each tank management method that contributed to the production of Category 4 (quality score >40; see Supplementary information) single-polyp microfragments used for cryopreservation experiments. **(C)** Quality scores for single-polyp microfragments across the total production period. **(D)** Single polyp microfragments ready for experimentation showing healthy, inflated, tissue and extended tentacles. **(E)** Proportions of single-polyp microfragments in each quality score category for each tank management method. Asterisk indicates a statistical difference within the same row (*alpha=*0.05).

Over the course of this study, three different approaches to tank management were utilised: ‘mini reef’ (MR), ‘clean start’ (CS), and ‘green film’ (GF). Although the GF method resulted in the highest proportion of coral colonies producing microfragments suitable for SPMF production (Fig. 2B), there was no significant difference between this approach and the MR (Fisher’s exact test, P = 0.1057) or CS (Fisher’s exact test, P = 0.0538) methods, possibly due to the low number of replicates in some tank methods. The development and use of these three approaches was an iterative process as our understanding of the biotic and abiotic conditions required to produce optimal coral material advanced, with the intended outcome being to maximise production of coral material suitable for cryopreservation. As such, the three methods were used consecutively so no direct, concurrent, comparisons between tank methods were possible. Over three thousand SPMFs were produced during the study, more than half of which were made during the period when the GF method was in use (Fig. 2C). In general, SPMFs were ready to use 48 to 72 hours after creation with each given a score based on observations of the health and integrity of the coral tissues and skeleton (see Supplementary information). Only SPMFs that received a score over 40 (Category 4) were deemed suitable for experimentation, which corresponded to SPMFs that had fully-healed inflated tissue with no signs of tissue loss or regression and a thin skeleton with no embedded skeletal debris (Fig. 2D). Fewer than 10% of the SPMFs produced were suitable for use in toxicity and cryopreservation experiments (Fig. 2E), with the remainder showing varying degrees of incomplete tissue or skeletal healing, and differences in the proportions of suitable SPMFs were observed between the three tank methods (Chi-square = 7.2492, df = 2, P = 0.0267). Although the proportion of Category 4 SPMFs produced using the CS tank method was significantly higher than both the MR (Chi-square pairwise nominal independence, P = 0.0236) or CS (Chi-square pairwise nominal independence, P = 0.0259) methods, the establishment and maintenance of the CS tank method was laborious and time consuming so was ultimately modified to create the less intensive GF approach. Although the GF method did not produce significantly more SPMFs than the MR method (Chi-square pairwise nominal independence, P = 0.6560), it was quicker to establish and resulted in a more consistent tank ecology that was easier to maintain.

### Vitrification solution development and testing

Development of a suitable vitrification solution took place over a nine-month period. The evolution of tank conditions, along with natural fluctuations in water temperature and differences in growth from colony-to-colony, meant that the supply and quality of coral material and subsequent robustness of SPMFs varied considerably over this time. Consequently, the response and recovery of SPMFs exposed to cryoprotectant solutions also varied, occasionally resulting in inconsistent or contradictory results from week-to-week. Initial assessments of toxicity and vitrification were based on the cryoprotectant solution developed for cryopreservation of coral larvae^11^. This solution showed low toxicity but was unable to vitrify consistently in droplets large enough to contain the SPMFs (2–4 µL). Development of a suitable vitrification solution that was both minimally-toxic and able to vitrify at the required volume was undertaken by assessing several complex solutions containing various combinations of cryoprotectants, ranging from approximately 3.2–6.0 M total solute content. Potential vitrification solutions that were able to vitrify in 4-µL droplets with no signs of haziness or cracking were then tested for toxicity to SPMFs. The final vitrification solution selected for toxicity consisted of 1M DMSO + 1M PG + 1M glycerol + 1.3M trehalose in PBS, which equated to approximately 4.6 M of total solutes or approximately 35% (w/w).

Refinement and assessments of cryoprotectant combinations were based on visual evaluation of tissue health and survival of SPMFs in response to exposure to the complex cryoprotectant solution. Immediately after exposure to cryoprotectant solutions SPMFs often showed signs of minor tissue damage (i.e., appearing ‘fluffy’) and increased mucous production, which typically resolved within 24 hours in surviving SPMFs. Assessment of SPMFs at 24–48 hours post-treatment revealed that exposure to vitrification solutions caused varying degrees of tissue damage (Fig. 3A–D) and in some cases death of the polyp (Fig. 3E–F). In most cases, surviving SPMFs showed signs of tissue remodelling (e.g., smoothing of damaged tissues, expulsion of skeletal debris and symbionts) within 48 hours of exposure to vitrification solution, and the presence of a clearly defined coral polyp was considered characteristic of survival. Tissue damage observed in surviving SPMFs included minor tissue regression and skeletal exposure at the edges (Fig. 3A–B), but some vitrification solution exposures caused more pronounced tissue regression and expulsion of symbionts (Fig. 3C), or, occasionally, complete loss of coenosarc tissue with only the polyp remaining intact (Fig. 3D). In some cases, dead SPMFs appeared to be relatively intact but on closer inspection showed a lack of tissue organisation (Fig. 3E), but most dead SPMFs showed obvious signs of tissue disintegration and lacked a clearly defined polyp (Fig. 3F).

**Figure 3.**
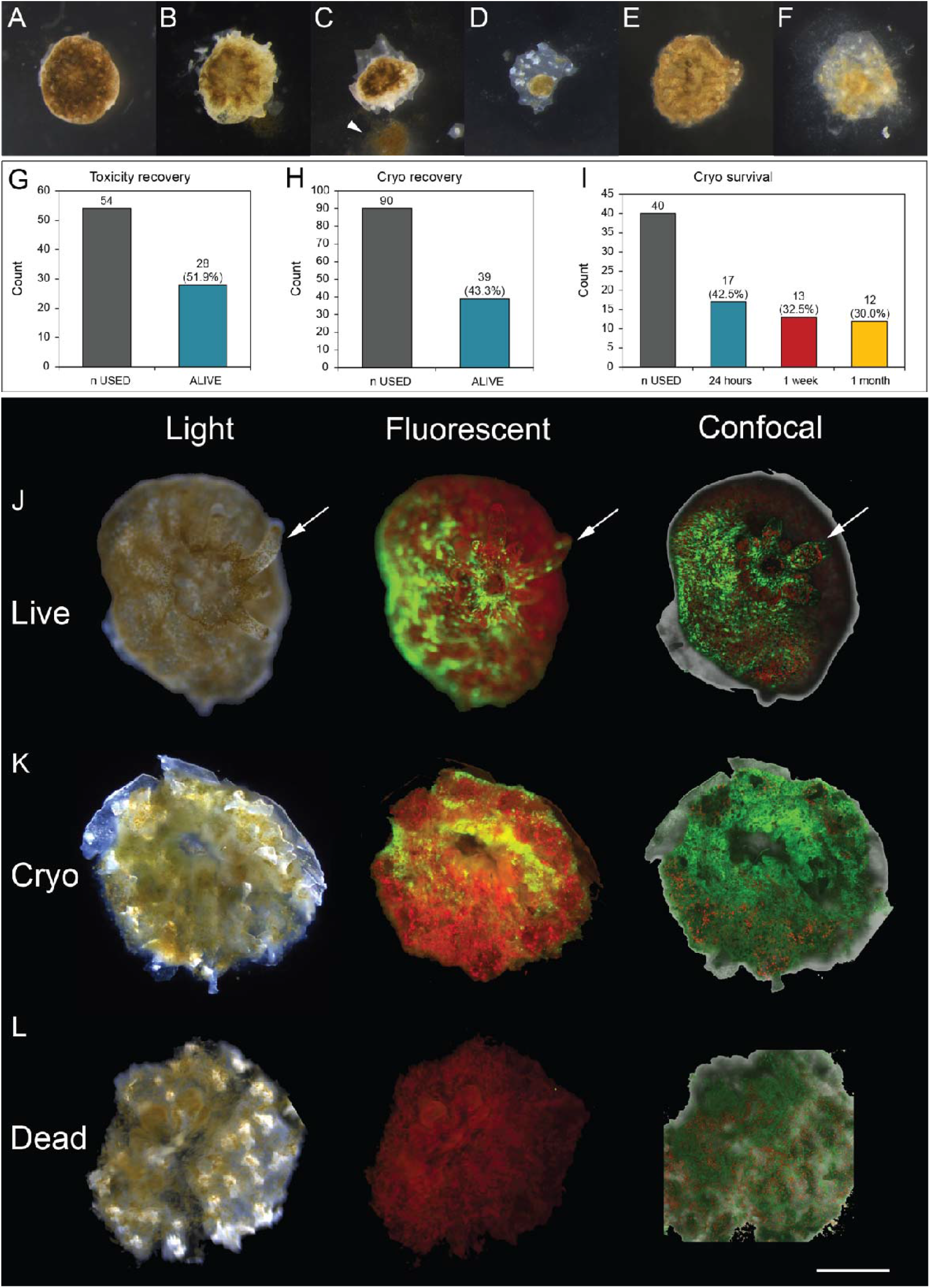
Response of single-polyp microfragments (SPMF) to toxicity and cryopreservation experiments. **(A–F)** Variation in tissue damage, including symbiont expulsion (arrowhead), and recovery observed 24 hours after exposure to vitrification solutions in surviving **(A–D)** and dead **(E–F)** single-polyp microfragment. **(G–I)** Total recovery counts of single-polyp microfragments at 24 hours post-treatment for toxicity **(G)** and cryopreservation **(H)** experiments over the course of the study and survival of a sub-set of vitrified and laser-warmed single-polyp microfragments followed for 1-month post-treatment in June–July 2021 **(I). (J–L)** Light, fluorescence, and confocal microscopy images of control untreated **(J)**, post-vitrification and warming **(K)**, and control dead **(L)** single-polyp microfragments showing characteristic tissue appearance and fluorescence. All SPMFs were produced using the GF tank method. Scale bar = 500µm.

Although a single-step one-minute exposure to vitrification solution was suitable for assessment of toxicity, rapid cooling of SPMFs exposed to vitrification solution in this way resulted in ice crystals in and around the mouth of the polyp, suggesting that this approach was insufficient for dehydration or cryoprotection. A longer, stepped, equilibration period was therefore developed to increase dehydration and potentially permit cryoprotectant permeation into tissues, which also necessitated the development of a more complex rehydration protocol. The equilibration process devised for the SPMFs consisted of a three-step dehydration and cryoprotectant loading process, followed by cryoprotectant unloading and rehydration over six-steps (Fig. 4). Exposure of SPMFs (n=54 SPMFs from 18 coral colonies; range 2–6 SPMFs/colony, mean 4.2 ± 0.49/colony) to the final vitrification solution using this protocol resulted in 51.9% mean survival (Fig. 3G). There was considerable variation (ranging from 0 to 100% survival) in the response of SPMFs to vitrification solution exposure from week-to-week, likely due to the colony-to-colony and environmental variability observed throughout the study. Toxicity experiments were conducted using the same batches of SPMFs as vitrification and laser nanowarming experiments, and on some occasions helped to explain poor post-warming outcomes or were used to determine whether vitrification and nanowarming experiments should proceed.

**Figure 4.**
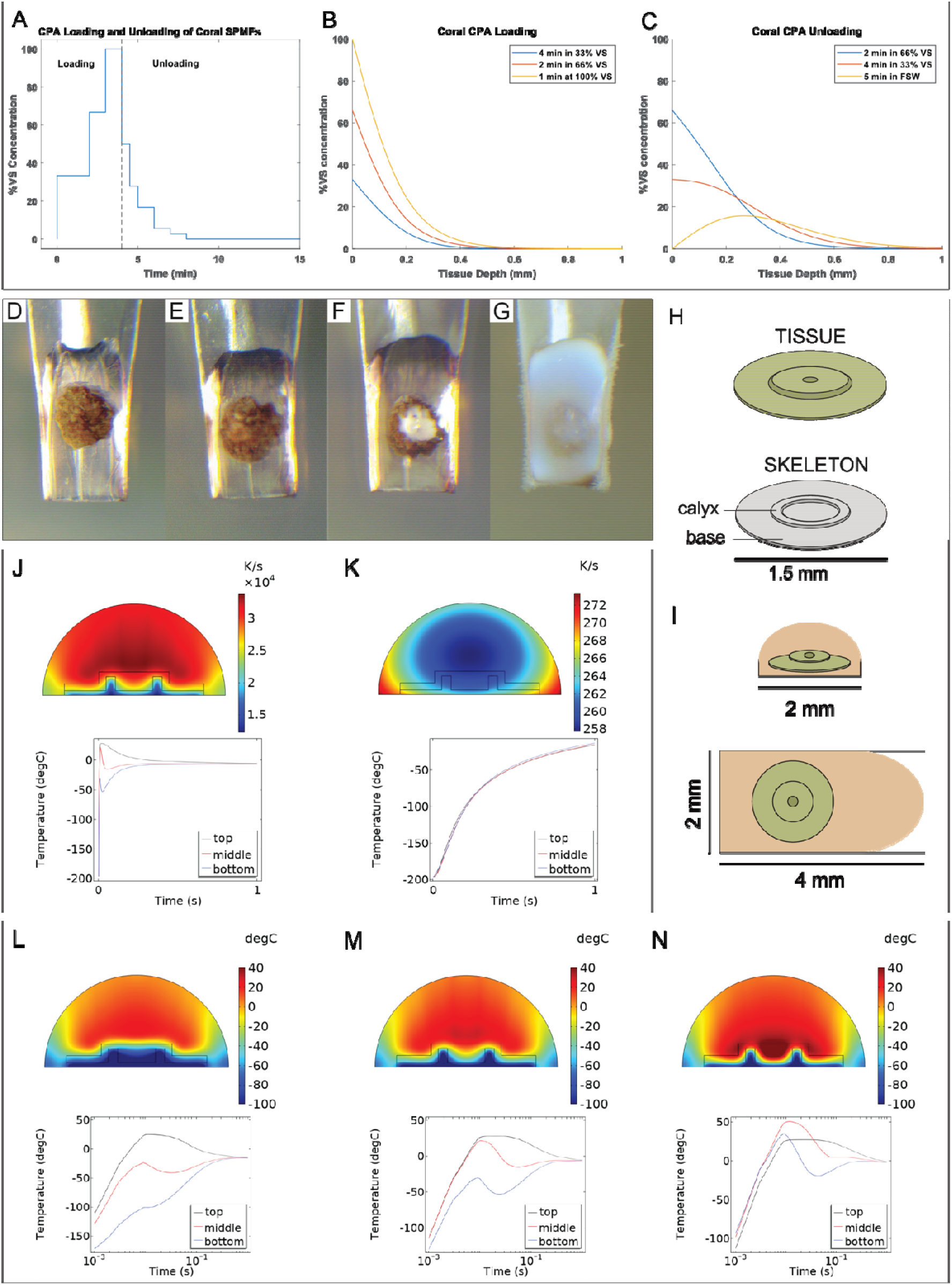
Vitrification and laser nanowarming of SPMFs from *Porites compressa*. (**A–C**) Equilibration and rehydration of single-polyp microfragments: (**A**) CPA step loading protocol with CPA concentration in tissue after each (**B**) loading step and (**C**) unloading step assuming a diffusion coefficient of 6 × 10^−11^ m^2^/s. (**D–G**) single-polyp microfragments were equilibrated and vitrified on modified cryotops: (**D**) single-polyp microfragment in vitrification solution (**E**) vitrified SPMF showing a clear glassy solid with no signs of ice formation; (**F**) single-polyp microfragment showing partial vitrification and ice crystal formation around the polyp mouth; (**G**) ice formation throughout the CPA droplet caused by devitrification during convective warming in air. (**H–I**) Simplified representation of the single-polyp microfragment and CPA droplet used to model warming profiles: (**H**) Coral skeleton consisting of a base and calyx that is overlaid by the coral tissue that covers the skeleton including the edges, with the polyp filling the calyx; (**I**) the single-polyp microfragment within the ∼2 × 4mm CPA droplet with a hemispherical profile on the cryotop. (**J–K**) Warming rate profiles in the droplet for (**J**) laser nanowarming (30,000 K/s) and (**K**) convective warming in 300K water (300 K/s), with plots corresponding to the temperature at the top, middle, and bottom of the coral single-polyp microfragment for laser nanowarming and convective warming respectively. (**L–M**) Temperature profiles immediately after laser pulse for coral single-polyp microfragments loaded with (**L**) no GNR, (**M**) 50% droplet GNR, and (**N**) 100% droplet GNR, with plots corresponding to the temperature at the top, middle, and bottom of the coral tissue for no GNR, 50% GNR, and 100% GNR respectively.

### Vitrification and laser nanowarming

Vitrification and laser nanowarming experiments took place over a ten-month period following the development of a suitable vitrification solution and protocols for cryoprotectant equilibration and rehydration. During this period, a total of 90 SPMFs from 15 coral colonies were vitrified and warmed (range 1–12 SPMFs/colony, mean 6.0 ± 1.0/colony), with an overall survival rate at 24 hours post-warming of 43.3% (Fig. 3H). Post-warming observations of a sub-set of SPMFs in June–July 2021 (Fig. 3I) found minimal decline in the survival rate after one week (32.5%) and one month (30.0%) compared to the initial survival at 24 hours (42.5%), indicating that the critical period for survival of SPMFs is the first 24 hours post-warming.

Fluorescence and confocal microscopy imaging of control (untreated) SPMFs showed that healthy coral tissues had well-organised green fluorescence, due to the presence of green fluorescent protein (GFP), while the algal symbionts appeared as red dots distributed throughout the tissues and the tentacles surrounding the polyp mouth (Fig. 3J). Immediately post-warming, SPMFs often showed signs of tissue damage like that observed in the toxicity experiments, with more pronounced damage apparent by 24 hours post-warming. Although the tissues surrounding the polyp mouth remained clear and intact in surviving SPMFs, the coenosarc showed signs of degradation and the symbionts in the tissues appeared a greenish colour instead of their normal brown, indicating that damage had occurred (Fig. 3K, Light). Fluorescence microscopy showed clearly defined GFP fluorescence around the polyp mouth post-warming; however, fluorescence was reduced in the coenosarc, and the tentacles were not visible (Fig. 3K, Fluorescence), indicating that the coral polyp was under stress. Similarly, confocal imaging showed clearly defined tissues around the polyp mouth but relatively few algal symbionts were present in the tissues (Fig. 3K, Confocal) compared to the untreated control. This diminished abundance of fluorescent symbionts post-warming, together with the colour change observed under light microscope, suggests poor survival of the symbionts following the vitrification and nanowarming process. The algal symbionts that live within coral are dinoflagellates from the Family Symbiodiniaceae and provide most of the energy requirements of their coral hosts via photosynthesis. These cells are known to have very different membrane permeability properties and cryopreservation requirements to coral tissues, and can be challenging to cryopreserve successfully ^20 21^. Moreover, the darker pigmentation of the symbionts may have enhanced their infrared laser absorption properties compared to the surrounding coral tissues and potentially impacted the survival of these cells.

The loss of algal symbionts and the photosynthetic energy that they provide to the corals may also have impaired the ability of the coral to heal damaged tissues post-warming, and there was generally less tissue remodelling observed post-warming SPMFs compared to SPMFs that were exposed to vitrification solution alone. By comparison, dead SPMFs showed significant degeneration of coenosarc tissues and lacked a clearly defined polyp (Fig. 3L, Light), and fluorescent and confocal imaging showed an almost complete loss of GFP fluorescence and disorganised distribution of algal symbionts in the tissues (Fig. 3L, Fluorescence and Confocal).

### Modelling of single-polyp microfragments

Cryoprotectant diffusion and thermal profiles were modelled to better understand the physical properties of the SPMF vitrification and laser nanowarming system and help to determine where the system may be improved in the future. The simulation results from the 1D planar diffusion model, shown as % cryoprotectant diffused as a function of tissue depth assuming a diffusion coefficient of 6 × 10^−11^ m^2^/s ^22^, indicated that the three-step cryoprotectant loading and six-step unloading protocol developed for the SPMFs (Fig. 4A–C) permitted the movement of cryoprotectants across coral tissues. The loading protocol resulted in 50% of the vitrification solution concentration (∼35% (w/w)) at a depth of 100 µm, which subsequently declined to under 20% at a depth of 200 µm. Given the diffusion coefficient of the CPA, and size of the SPMF, we expect CPA to continue diffusing throughout the specimen until vitrification. The diffusion of cryoprotectants into the coral tissues permitted sample vitrification with no visible ice crystals (Fig. 4E), with vitrified SPMFs appearing similar to those in vitrification solution prior to immersion in liquid nitrogen (Fig. 4D). By contrast, incomplete cryoprotectant loading resulted in ice crystal formation in and around the polyp mouth where the coral tissues were at their thickest and most complex (Fig. 4F), while extensive ice crystal formation was observed throughout the sample when SPMFs were immersed in liquid nitrogen without vitrification solution or allowed to warm convectively in air (Fig. 4G). The warming rates generated throughout the coral using laser nanowarming were 100 times faster than those generated from convective warming (Fig. 4J–K), indicating that the laser nanowarming method can still allow for successful cryopreservation using much lower concentrations of CPA than convective methods.

A major factor in the success of laser nanowarming is the distribution of gold nanoparticles within and surrounding the tissues. The incubation of SPMFs in GNRs in FSW for an hour prior to vitrification was designed to permit passive diffusion of the GNRs into the coral tissues. As it was not possible to determine the exact extent of GNR permeation into coral tissues during this incubation period, warming simulations were based on theoretical concentrations of GNR within the tissues of 0, 50, and 100% of the surrounding vitrification solution (Fig. L–M). The warming rates for the 0%, 50% and 100% examples were 15,000, 30,000, and 50,000 K/s respectively. These simulations indicated that with a modest uptake of 50% of droplet GNR within the SPMF we can expect to exceed the critical warming rate for vitrification solutions at total solute concentrations above 35% (w/w) (Fig. 4M). For SPMFs thicker than 100 µm, warming rates in the tissue need to be substantially higher than 30,000 K/s to avoid ice crystallization. Since increasing the CPA loading times would likely increase tissue damage and death from toxicity, our results show that thinner fragments are perhaps best suited for cryopreservation. The fact that living tissues were observed post-warming suggests that there was some passive diffusion of GNRs into the tissues during the incubation step, although it was likely less than the 50% of external GNR concentration used for modelling. To increase the concentration of GNR within the tissue it may be necessary to devise methods for microinjections of GNR and CPA into the polyp mouth or increase the duration of incubation of SPMFs in GNR, to improve post-warming recovery.

## Conclusions

This study describes a system for cryopreserving adult coral tissues for the first time, utilising a novel sample type, single-polyp microfragments, suitable for vitrification and nanowarming. Adapted and applied to other reef-building species, SPMFs may be a useful sample type for screening experiments that require large numbers of treatments and coral replicates, for example chemical toxicity assays or testing the efficacy of antibiotic or supplement treatments to aid recovery from bleaching or disease. The advances made, in particular the development of a vitrification solution and cryoprotectant loading and unloading protocols, can form a basis for future development of a field-ready protocol for cryopreservation of adult coral tissues, enabling biobanking of threatened coral species throughout the year. Key to this approach will be the refinement of methods to nurse coral through the first few days post-warming and getting them to a point where they can re-uptake algal symbionts into their tissues to replace those lost during cryopreservation. The loss of symbionts during the vitrification and laser nanowarming process likely impacted the ability of the SPMFs to heal and recover, highlighting the importance of cryopreserving and banking coral symbionts alongside coral cells and tissues to enable replacement of damaged symbionts as part of recovery post-warming. The rapid advancement of cryopreservation technologies by programs such as the U.S. National Science Foundation Engineering Research Center ATP-Bio (www.atp-bio.org) means that new approaches may soon be available to overcome these issues. By building on the work in the present study, these technologies may someday permit cryopreservation of larger microfragments with multiple polyps to support biobanking of threatened coral species for reef restoration and adaptation activities.

## Methods

### Colony Collection and Care

*Porites compressa* colonies were collected from various reefs in Kaneohe Bay, Oahu, HI in accordance with Special Activity Permits (permit # 2020-25, 2021-33, and 2022-22) from the Department of Land and Natural Resources of the State of Hawaii. Care was taken to collect colonies at different locations on each reef to avoid collecting clones of the same genotype. Collected colonies were housed outdoors in fiberglass tanks (92 × 61 × 46 cm) fed with filtered seawater at a rate of ∼1.5L/min and supplied with 2 × 4 cm airstones for water circulation (Pentair, Aquatic Ecosystems, Apopka, FL). Seawater was filtered by 100-, 50-, and 25-µm bag filters, and a 24” sand filter (Hayward Industries Inc, Newport News, VA) to remove detritus, algae, and pest invertebrates.

Corals received natural sunlight, which was attenuated to reduce intensity via an overhead, metal frame tent structure covered with 50% attenuation shade cloth (Magna Distributors Inc., Honolulu HI), and by individual tank lids, constructed of a PVC frame covered with a clear polycarbonate sheet, to serve as a rain guard (Palram Americas, Kutztown, PA). During summer, clear polycarbonate was replaced with tinted polycarbonate (Palram Americas, Kutztown, PA), reducing incoming light by a further 30%, and each rain guard was half-covered with an additional section of 50% shade cloth. This shading scheme resulted in tanks experiencing Photosynthetic Active Radiation (PAR) values between 200-1500 µmols/m^2^/sec during daylight hours measured by an electronic PAR meter (Apogee Instruments, North Logan, UT).

### Colony Microfragmentation

Coral colonies were progressively fragmented using a diamond bit, rotary cutting tool (Robert Bausch Tool Corporation, Mt Prospect, IL) then cut into flat 0.5 cm strips of tissue using a diamond band saw (www.gryphoncorp.com) to yield thin 0.5–1.0 cm sections of tissue called microfragments^12^. Microfragments were glued to a half sheet of heavy duty, plastic paper protector (Samsill Corporation, Ft Worth, TX) using a drop of extra thick super glue gel (www.bulkreefsupply.com), spaced apart such that 30 fragments of 2 genotypes were mounted to each sheet. Paper protectors were clipped to 0.25 cm thick, clear acrylic sheets of equal size (Plaskolite, Columbus, OH) with plastic seaweed clips (Amazon.com) for easy manipulation and excision of fragments and maintained in flow-through seawater tanks for grow out.

### Microfragment Care

Microfragment grow-out tanks were setup identically to colony holding tanks. During the Winter months, temperatures were maintained between 25 and 26.5°C using 2 × 500 W titanium heaters (finnex.net) per tank controlled by an external controller (Inkbird Technical Corporation Limited, Shenzen, China). Controllers were programmed such that temperature did not fluctuate by more than 1.3 °C in a day, confirmed using a bluetooth temperature logger that recorded temperature every 30 seconds (Inkbird Technical Corporation Limited, Shenzen, China). Salinity and pH were checked daily with handheld meters (Sper Scientific, Scottsdale, AZ; and Milwaukee Instruments Rocky Mount, NC) to ensure minimal fluctuation in abiotic conditions. During Summer months, grow-out tanks received filtered seawater from a sump cooled to 25.5°C to ensure that water temperature in the grow-out tanks remained below 28ºC. Water in the sump was cooled using a recirculating 1.5 HP chiller (ecoplususa.com) with an internal 4800 L/h water pump (www.ecoplususa.com) and was pumped to each grow-out tank with a 5815 L/h external water pump (Iwaki America, Holliston, MA). All chilled water reaching grow out tanks was flow-through and was not recirculated to the sump to prevent transfer of biofouling organisms between tanks.

### Control of Biotic Factors

Tank ecology was maintained as described in Page et al. ^19^. Briefly, prior to the introduction of microfragment plates, the ecology of a new grow-out tank was established by seeding the tank immediately upon start-up with 10 individual 2 cm chips of crustose coralline algae (CCA), and 20 *Trochus intextus* snails were added to control fast growing algae. After 1 week in the tank, the tank was scrubbed thoroughly to remove fast-growing diatoms, then drained, pressure washed with filtered seawater, and rapidly refilled, ensuring that the sidewalls always remained damp. This process was repeated weekly until signs of green film and CCA recruitment were present on the tank’s sidewalls, which generally took about 2 weeks in our system.

Once beneficial tank ecology was established, newly cut microfragment plates were added to the tank and the number of *Trochus* was increased to 60-80 individuals, with 40 snails per tank allowed to graze freely and the remainder divided evenly between 5 inverted 15–25-cm baskets held with magnets to the walls and positioned over dark film algae. Each morning all baskets were re-positioned, and each tank was siphoned of detritus. When in use, sump tanks underwent the same tank ecology selection process as grow out tanks.

### Single-polyp microfragment production

Microfragments were selected for use based on the amount and thickness of sheeted growth, development of the polyps, and overall health of the microfragment. Suitable microfragments had multiple rows of clearly defined coral polyps (i.e., with fully developed mouth and tentacles), with thin skeleton showing no signs of calyx development, and even distribution of symbionts throughout the tissues with no signs of tissue loss or damage. Selected microfragments were excised from the growing plate and placed into individual wells of a 6-well plate containing tank seawater. Well plates were brought to the lab and placed in an incubator (Percival Scientific, Indiana, USA) set at 26ºC, with 80–110 µmols/m^2^/sec PAR for up to four hours prior to creation of SPMFs.

For creation of SPMFs, individual microfragments were gently rinsed with seawater using a transfer pipette and moved to a 70-mm dish containing 0.22-µm filtered seawater (FSW) on the stage of a dissecting microscope (Olympus SZX12). Individual polyps were excised from the sheeted area of the microfragment using a 1.5-mm biopsy punch (Cat. # RBP-15, Robbins Instruments, USA) and collected in a second 70-mm dish by gently flushing FSW through the biopsy punch using a transfer pipette. Single-polyp microfragments were cleaned to remove skeletal debris and mucous by gently pipetting the SPMF back and forth with a cut pipette tip or transfer pipette, then transferred to a clean 6-well plate containing FSW (6 SPMFs per well containing 10 mL FSW) and maintained in an incubator. Handling of SPMFs throughout production and experimentation was achieved using a P200 micropipette set to 50 µL, with the attached pipette tip trimmed to increase the diameter of the opening. Single-polyp microfragments were drawn gently into the tip with 50 µL of FSW and allowed to settle towards the tip opening by holding the pipette upright. The SPMF was released by gently touching the pipette tip to the surface of the FSW without expelling any of the water in the tip, allowing the SPMF to drop into the new well. Single-polyp microfragments were viewed under a dissecting microscope and flushed gently with the pipette if necessary to ensure that the polyp was upright. This method permitted gentle handling and movement of the SPMFs with minimal transfer of liquid between steps. At 24-h intervals following production, SPMFs were transferred into new 6-well plates with clean FSW and assessed using metrics designed specifically to track healing and suitability for experimentation (see Supplementary information). Single-polyp microfragments were produced on a weekly cycle, depending on the availability of suitable microfragment material, which typically resulted in around 10–20 SPMFs from one to three coral genotypes available for experimentation in any given week.

### Vitrification solution toxicity assessment

Development of a suitable vitrification solution was undertaken by assessing the toxicity and ability to vitrify of a range of vitrification solutions (n = 15) containing various combinations of dimethyl sulfoxide (DMSO, Cat. #D4540, Sigma Aldrich), 1,2-propanediol (PG, Cat. #398039, Sigma Aldrich), ethylene glycol (EG; Cat. #324558, Sigma Aldrich), polyethylene glycol (PEG, Cat. #807485 Sigma Aldrich), glycerol (Cat. #G7893, Sigma Aldrich, and trehalose (Cat. #T-104-4, Pfanstiehl) at total solute concentrations ranging from approximately 3.2 to 6.0 M of total solutes.

Vitrification solutions were first assessed for their ability to vitrify by placing a 4µL droplet of the solution onto a modified version of the Kitazato CryoTop (Kitazato, Tokyo, Japan) like the one described previously for larvae^11^, consisting of a 2 × 10 mm polypropylene blade mounted in a wooden press-fit holder. The cryotop was then plunged into liquid nitrogen the droplet visually inspected for signs of ice crystal formation (i.e., haziness within the droplet) or cracking. Solutions that were able to vitrify were then assessed for toxicity by exposing individual SPMFs to the vitrification solution for one minute, followed by a single-step rehydration in 10% DMSO in FSW for one minute, then returning the SPMF to FSW in a 24-well plate in the incubator for recovery. To do this, SPMFs were picked up with a pipette as described and transferred to a 35-mm dish in a 50-µL droplet of FSW. Seawater was removed with a pipette, and a 10 µL droplet of vitrification solution was placed onto the SPMF and mixed by pipetting gently back and forth. After one minute, the SPMF was picked up with a pipette and transferred to a 35-mm dish containing 10% DMSO in FSW, with gentle pipetting to rinse residual vitrification solution from the polyp. After one minute, the SPMF was transferred to the well of a 24-well plate containing 2 mL of FSW. This process was repeated for at least three SPMFs per vitrification solution. Well plates containing treated SPMFs were placed in an incubator set to 26ºC, with 80–110 µmols/m2/sec on a 10:14 light:dark cycle, and SPMFs were imaged immediately after treatment and then at 24 h, 48 h, and one week after treatment to assess recovery. Assessment of treated SPMFs was based on tissue integrity, symbiont density / tissue colour, and the morphology of the mouth and tentacles (Fig. 3).

For refinement of the equilibration and rehydration protocols, exposure to stepped increases and decreases in vitrification solution concentration were performed using a modified cryotop. Individual SPMFs were placed onto the cryoblade with the holder positioned on the stage of a dissecting microscope, ensuring that the polyp was upright. Excess seawater was removed with a pipette and the cryoblade blotted with a Kimwipe. A 5µL droplet of 33% vitrification solution (1:2 dilution of vitrification solution with FSW) was placed onto the SPMF and mixed by pipetting back and forth to rinse off residual FSW and ensure vitrification solution contact with the tissues. After 2 min exposure the 33% solution was pipetted off, the cryotop carefully blotted with a Kimwipe to remove excess solution, and a 5µL droplet of 66% vitrification solution (2:1 dilution of vitrification solution with FSW) placed onto the SPMF and mixed by pipetting. After 1 min, the 66% solution was removed, excess blotted away, and a 4µL droplet of full-strength VS was added with gentle pipette mixing. After one minute exposure to the full-strength vitrification solution the SPMF was rehydrated by moving it through a timed series of solutions with decreasing osmolality (Fig. 4). The first rehydration step consisted of 100µL of 50% vitrification solution (1:1 dilution of vitrification solution with FSW) with a calculated osmolality of approximately 2800 mOsm, which was used to rinse the SPMF from the cryoblade into a 35-mm dish, ensuring that the rehydration solution stayed as a discrete droplet surrounding the SPMF. Subsequent osmotic steps (2000, 1600, 1200, and 1100 mOsm) were achieved by adding FSW to this droplet for the required time, as described in Fig. 4. After the final step the SPMF was transferred to a 24-well plate in 2 mL of FSW, which was kept in the incubator. Single-polyp microfragments were imaged immediately after treatment and then at 24 h, 48 h, and one week to assess recovery using an Olympus SZX12 dissecting microscope, with an Infinity 3S Lumenera camera and Infinity Analyze software for visualization.

### Vitrification and laser nanowarming

Vitrification and laser nanowarming of SPMFs were carried out on a modified cryo blade, made by pinching a 4 × 10 mm strip of polypropylene (C-Line Products Mount Prospect, IL) around the point of a pair of fine-point forceps to create a narrow channel approximately 2 mm wide with 1mm high walls on each side. This channel cryo blade was then mounted in a wooden press-fit holder and handled as described. The channel cryo blade was devised as a means of maintaining a relatively even layer of vitrification solution over the tissue surface of the SPMF to minimise lensing of the infrared laser pulse and more evenly warm across the diameter of the SPMF. Healthy Category 4 SPMFs were selected using an Olympus SZX12 dissecting microscope and transferred into a 35-mm dish containing FSW with 0.07 mg/mL (4.5 × 10^11^ np/m^-3^)1064nm resonant PEG-coated gold nanorods ^14^ (GNR; NanoComposix, San Diego, USA) for one hour. After the pre-treatment period, SPMFs were placed on the channel cryotop blade individually, excess water was removed with pipette and the cryo blade blotted with a Kimwipe. The SPMF was equilibrated with cryoprotectants using the 33% and 66% vitrification steps described above, followed by a 4µL droplet of full-strength VS containing 0.07 mg/mL GNR. To help ensure the consistency of sample melting, GNRs were vortexed and sonicated prior to mixing with the vitrification solution, and this solution was used within an hour of preparation. The wooden holder with the cryo blade and SPMF was then transferred into the chamber of a bench-top iWeld 980 Series, 60 joule, Nd:YAG unpolarised infrared laser (model number 585-986-080; LaserStar Technologies Corporation, Orlando, FL, USA) and inserted into the arm of a custom-made cryo laser apparatus (Design Solutions, Inc., Chanhassen, MN, USA) for vitrification and laser nanowarming as described in Daly et al.^11^. After 1 min exposure to the full-strength vitrification solution, the cryo apparatus was used to lower the sample into a liquid nitrogen bath for vitrification. The sample remained in the liquid nitrogen for 10 seconds to ensure temperature equilibration, then the cryo apparatus was used to raise it into the laser focal region. The sample was held briefly (<3s) in this position to allow excess liquid nitrogen to evaporate and permit assessment of vitrification, then warmed with a single pulse from the infrared laser (240V, 9.0ms, ∼ 8.2 J) using the iWeld Spike Profile setting. The vitrification and nanowarming procedure was observed through the iWeld microscope headpiece to monitor for signs of ice crystal formation or devitrification, which typically appeared as increased opacity or cracking of the droplet or SPMF. If signs of ice were observed in the tissues, then the SPMF was discarded, while those that remained clear were rehydrated as described above and transferred to individual wells in a 24-well plate containing 2 mL FSW. At the end of the experiment the well plate was placed into the incubator for recovery, with foil covering the lid for the first 24h post-treatment to reduce light stress to the symbionts. Post-thaw SPMFs were imaged at 24 h, 48 h, and one-week post-treatment to assess tissue integrity, symbiont density / tissue colour, and the morphology of the mouth and tentacles as described previously and assigned a score from 1 to 4, with scores of 1–3 corresponding to live SPMFs with increasing levels of tissue damage and 4 corresponding to those that were dead. Additionally, confocal imaging of the SPMFs at 24h was used to identify GFP autofluorescence as a marker of tissue survival.

Laser scanner confocal microscopy (LSCM) can visualise coral auto-fluorescent proteins as well as coral algal symbionts, allowing for a non-invasive assessment of both coral tissue and symbiont health ^23^. Samples were scanned and imaged using Zeiss LSCM 710 and Zen Black processing software (2011 v14.0.16.201). Excitation wavelength of 405 nm with a Diode laser (UV) allowed for visualization of coral fluorescent proteins and algal symbiont fluorescence at a magnification of 10x. Fluorescent protein fluorescence was collected 455-620nm, and symbiont fluorescence was collected at 620-720nm. Single-polyp microfragments were imaged in a glass bottom dish (0.17 mm) filled with filtered seawater for inverted imaging. Scanned z-stack slices were 10 µm apart and integrated using maximum intensity projection.

### Simulating CPA diffusion and photothermal heat transfer in single-polyp microfragments

To estimate the laser absorption and heat generation, Monte Carlo simulations for light transport within the droplet and coral SPMF were performed using methods developed previously^16, 24^. Briefly, the coral SPMF skeleton was modelled as a disk (base) and ring (calyx), sitting atop each other as shown in Figure 4H. The base had a diameter of 1.5 mm and thickness of 50 µm while the calyx had a diameter of 500 µm and height of 200 µm. Coral tissue filled the ring cavity and sat atop the rest of the skeletal structure with a varying thickness of 50 to 100 µm. The surrounding droplet was modelled as a hemisphere with a diameter of 2mm Figure 4I. Thermophysical and optical properties of the coral skeleton were modelled after those of human bone while those for the coral tissue were modelled after tissue perfused with PBS+2M glycerol. A table summarizing the properties used in the model can be found in the Supplementary information. Monte Carlo simulations to estimate the heat generation were carried out for varying amounts of GNR in the tissue to model the worst, realistic, and best cases for warming corresponding to a GNR concentration of 0%, 50%, and 100% inside the tissue relative to the droplet GNR concentration. The Monte Carlo simulation results were then analysed in COMSOL to generate the predicted temperature profiles in the droplet and SPMF given the laser settings (240V, 9.0ms, 2mm beam diameter) and droplet GNR concentration of 0.07 mg/mL. We estimated CPA uptake within the coral SPMF using 1D planar diffusion model with an assumed coral tissue diffusivity of 6 × 10^−11^ m^2^/s ^22^. Since no data was available for diffusion coefficient of coral tissue, estimates were made based on previous measurements of cardiac tissue of similar thickness.

## Statistical analyses

Data on microfragment production by tank method were analysed by Fisher’s exact test of independence and pairwise nominal independence analysis with ‘fdr’ adjustment, and single-polyp microfragment production data were analysed by Pearson’s Chi-square test and pairwise nominal independence analysis with ‘fdr’ adjustment, using R (R Core Team 2019) with the “rcompanion” (Mangiafico 2019) package. Data on recovery of SPMFs following toxicity and vitrification and laser nanowarming experiments were analysed in Excel, and are expressed as count or percentage data, with means expressed as ± SEM.

## Acknowledgements

This work was supported by Smithsonian Institution, The Smithsonian’s Women’s Committee, the Paul M. Angell Family Foundation, OceanKind, Revive & Restore, the Scintilla Foundation, the Zegar Family Foundation, the William H. Donner Family Foundation, the Cedar Hill Foundation, and the Hawaii Institute of Marine Biology. National Science Foundation Grant No. EEC 1941543 supported the Engineering Research Center Advanced Technologies for the Preservation of Biological Systems (ATP-Bio) is gratefully acknowledged. Coral collections were performed with the appropriate permits from the state of Hawaii’s Department of Land and Natural Resources (Special Activity Permit # 2019: 2012-63, 2020: 2013-47 and 2021: 2015-17). No institutional ethical approval was required for any of the experimental research described herein; however, animals were maintained with the highest husbandry standards. This manuscript has contribution number XXXX from the Hawaii Institute of Marine Biology.

## Author contributions

JD, JB, RP, CP, CL, and KH conducted the experimental work. KK and JK performed the modelling analyses. MH and JCB supervised the research. JD analysed the data, prepared the figures, and wrote the manuscript. All authors contributed to manuscript review and editing.

## Data availability

Data are available on request.

