## Supplemental Material for "The first proof of concept demonstration of nanowarming in coral tissue"

### Nanowarming of coral – Supplementary information

**Assessment of single-polyp microfragment healing:**

Photos were taken using an Olympus SZX12 dissecting microscope, Infinity 3S Lumenera camera, and Infinity Analyze for visualization. Single-polyp microfragments (SPMFs) were photographed the day they were cut, at 48 hours, and at 1 week. At 48 hours and 1 week, they underwent a metric assessment to determine the quality and health of each individual nanofragment. The three metrics assessed were 1) tissue coverage: the percent of healthy tissue surrounding the polyp and to the edge of the nanofragment; 2) tissue integrity: the amount of tissue regression and health of the tissue; 3) skeleton: the size of the excess shards of skeleton that the SPMF is expelling and the health of the skeleton. Each metric was given a value, as defined below.

To be suitable for cryoprotectant toxicity or vitrification and laser nanowarming experiments SPMFs needed to meet a minimum score for each metric, which was set at a score of 4 or 5 for Tissue coverage, 5 for Tissue integrity, and 4 for Skeleton. For each of the three metrics, an additional 10 points were added if these minimum requirements are met, for a maximum of 30 additional points in total. This was done to ensure that suitable SPMFs were clearly separated from those that scored highly in some metrics and poorly in others. Single-polyp microfragments were thus separated into 4 categories: Category 1 = score 0-10, Category 2 = score 11-29, Category 3 = score 30-39, and Category 4 = score 40-44, with only those SPMFs in Category 4 meeting the quality required for experimentation.

**(1) Tissue coverage (to the edge of the nanofragment)**

5 100% coverage with inflation

4 100% coverage without inflation

3 75% coverage

2 50% coverage

1 25% coverage

0 0% coverage

**(2) Tissue integrity (regression and health)**

5 0% regression-freshly made nanofrags and/or good looking nanofrags

4 25% regression

3 50% regression

2 75% regression -just the polyp could be present

1 100% regression -just the polyp could be present

0 Tissue disintegration

**(3) Skeleton**

4 No shards present around the nanofragment

3 Small shards present

2 Big shards present

1 Reduction of skeleton with or without shards

0 Disintegrated skeleton

| **Properties** | **Compartment** | **Value** | **Reference** |
| --- | --- | --- | --- |
| Density | Skeleton | 2.94 g cm^-3^ | ^1^ |
| Thermal Conductivity |  | 0.58 W m^-1^ K^-1^ | ^2^ |
| Specific Heat |  | 1260 J kg^-1^ K^-1^ | ^3^ |
| Absorption Coefficient |  | 0.2 cm^-1^ | ^4^ |
| Scattering Coefficient |  | 20 cm^-1^ |  |
| Density | Tissue + CPA | 1.00 g cm^2^ | water density |
| Thermal Conductivity |  | PBS+2M glycerol* | ^5^ |
| Specific Heat |  | PBS+2M glycerol ref* |  |
| Absorption Coefficient |  | 0.5 cm^-1^ | ^4^ |
| Scattering Coefficient |  | 10 cm^-1^ |  |
| Scattering Anisotropy |  | 0.94 |  |

Table (S1). Thermophysical and optical properties used in Monte Carlo light transport simulations and COMSOL Multiphysics FEM heat transfer analysis.
